# ATGL-catalyzed lipid catabolism promotes DNA double-strand break repair

**DOI:** 10.64898/2026.04.03.716381

**Authors:** Mahima Devarajan, Rachel K. Meyer, Gavin Fredrickson, William A. Hofstadter, Mara T. Mashek, Mari V. Reid, Evan W. Kerr, Anna Bartelt, Peter Gutenberger, Simone Intriago Tito, Linshan Laux, Emma Petta, Hai Dang Nguyen, Douglas G. Mashek

## Abstract

An imbalance of DNA damage over DNA repair contributes to the genomic instability that drives aging and numerous age-related diseases. While numerous DNA repair mechanisms have been elucidated over decades of study, little is known about the contribution of metabolism to genomic stability. We report that adipose triglyceride lipase (ATGL), a lipolytic enzyme, promotes DNA repair. We show that lipid droplets (LDs) accumulate in response to DNA damage and that inhibition of LD biogenesis before genotoxic stress increases the persistence of DNA damage. Overexpression of ATGL, which increases lipolysis, reduces DNA damage following etoposide and ionizing radiation, thereby promoting genomic stability. Further, ATGL expression prior to DNA damage attenuates the long-term consequences of DNA damage, reducing senescence and enhancing viability. Mechanistically, ATGL promotes double-strand break repair via NHEJ and HR to mitigate DNA damage. Overall, these studies reveal a novel role for LDs and their proteins in DNA damage and repair, thereby unveiling a mechanism by which lipid metabolism contributes to genomic stability.

## INTRODUCTION

DNA damage occurs thousands of times per cell per day, occurring from both exogenous (e.g., ionizing radiation, chemotherapeutic drugs) and endogenous (e.g., reactive oxygen species, transcription/replication conflicts) sources ^1,2^. While cells have evolved numerous mechanisms to recognize and respond to genotoxicity, these are not always sufficient for complete DNA repair, leading to several possible fates. First, persistent DNA damage can lead to permanent cell cycle arrest or senescence at the cellular level and premature aging at the physiological level ^3^. Next, incorrect or incomplete DNA repair can lead to acquired mutations and malignant transformation ^1,4^. Finally, DNA damage that is too severe for repair will induce apoptosis, resulting in tissue damage and loss of function. Therefore, the balance between DNA damage and the evolved endogenous mechanisms of DNA repair ultimately dictates genomic integrity.

Upon DNA damage, numerous signaling cascades are initiated to recognize and respond. One of the earliest events in response to double-strand breaks (DSBs) is the phosphorylation of serine 139 on histone H2Ax (commonly denoted γH2AX), which amplifies the DNA damage response and recruits DSB repair proteins ^5,6^. Indicative of the critical role of γH2Ax in DNA repair, mice lacking histone H2Ax show chromosomal aberrations, higher radiation sensitivity, and a failure to recruit repair proteins to break sites ^7^. Cells also initiate the critical shift from growth and proliferation to cell cycle arrest and DNA repair. Cells primarily repair DSBs with nonhomologous end joining (NHEJ) or homologous recombination (HR). NHEJ is considered the predominant form of DNA repair, involving the direct ligation of broken DNA ends and occurring at all stages of the cell cycle ^8^. In contrast, HR is characterized by its ability to utilize a homologous DNA sequence and is therefore often considered error-free ^9,10^. Despite decades of accumulated knowledge on DNA damage and DNA repair, DNA damage-associated diseases (cancer, age/senescence-related pathologies, etc.) remain persistent. Novel perspectives on the cell’s response to DNA damage are essential for further developing approaches to mitigate the widespread consequences of genomic instability.

Cytosolic lipid droplets (LDs) serve as hubs of lipid storage, metabolism, and signaling ^11^. Though commonly viewed for their roles in energy storage within adipose tissue, LDs accumulate in all cell types and play numerous roles in cell signaling and metabolism. Adipocyte triglyceride lipase (ATGL, gene name is PNPLA2) catalyzes the first and rate-limiting step of lipolysis, in which triglycerides are hydrolyzed to generate diacylglycerol and free fatty acids. ATGL preferentially channels hydrolyzed fatty acids to oxidative or signaling pathways ^12,13^. Indicative of its physiologic importance, ATGL knockout mice show impaired stimulated lipolysis and reduced fasting serum fatty acid levels, accompanied by increased adipose depot size and profound LD accumulation in the numerous tissues including the heart and liver ^14^. Tissue-specific knockouts of ATGL reveal a significant contribution to lipid catabolism in most cell types studied, highlighting a role beyond adipose tissue ^15^. ATGL has emerged as a key signaling hub, regulating numerous downstream nutrient-sensing and/or transcriptional networks that govern diverse biological processes ^16,17^.

In this work, we explored the connection between LD metabolism and DNA damage, highlighting a robust role for ATGL in promoting genomic stability. We show that lipid catabolism can promote DNA repair, prevent the induction of senescence, and maintain cell viability in response to genotoxic stress, thereby identifying new mechanisms that underlie the metabolic regulation of DNA damage and repair.

## RESULTS

### LD ablation prior to genotoxic stress increases DNA damage

We first sought to test if LD metabolism was altered by genotoxic stress. To interrogate LD dynamics after DNA damage, we subjected multiple cell lines to different forms of DNA damage and measured LDs by staining with BODIPY 493/503. AML12 cells and primary MEFs, mouse hepatocytes and fibroblasts, respectively, accumulated LDs with etoposide (a topoisomerase II inhibitor and a DSB inducer) in a time-dependent manner (**Fig 1A**). Primary MEFs and human IMR90s are both primary cell lines capable of replicative senescence via accumulation of DNA damage with repeated passages. These cells also accumulate LDs with replicative stress, indicating multiple types of DNA damage induce LDs (**Fig 1B**). Consistent with the literature, these data show that multiple cell types of human and mouse origin accumulate LDs in the presence of DNA damage ^18^.

**Figure 1.**
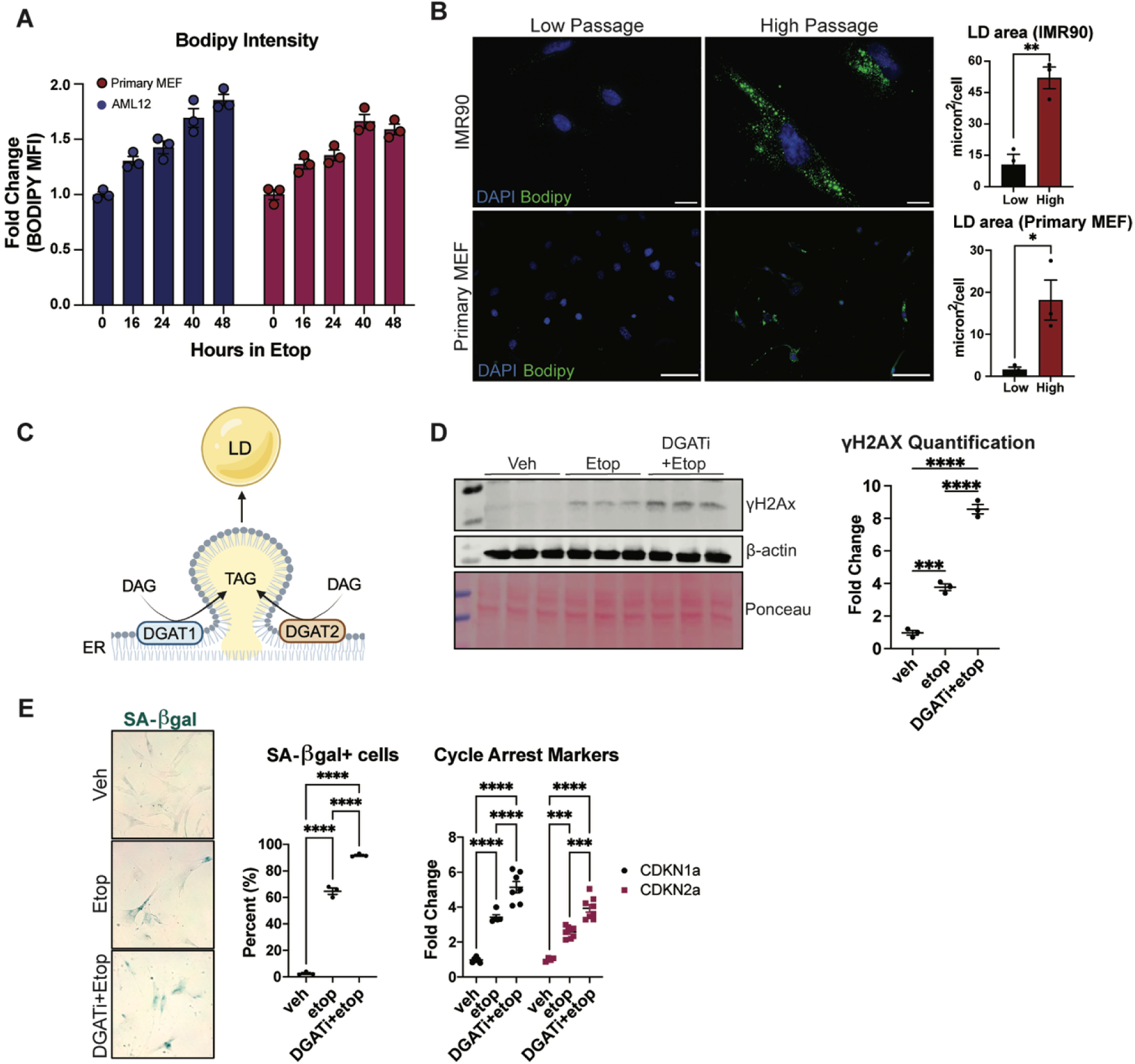
DGAT1/2 inhibition prior to genotoxic stress increases DNA damage. **A)** AML12 cells and primary MEFs were treated with 30 µM etoposide and harvested at the denoted timepoints of etoposide treatment. LDs were stained with BODIPY, and mean fluorescence intensity (MFI) was measured via flow cytometry. **B)** Primary MEFs and IMR90s were repeatedly passaged (Primary MEFs, low passage = P2, high passage = P6; IMR90, low passage = P6, high passage = P22. Primary MEF scale bar = 100 microns. IMR90 scale bar = 20 microns. A minimum of 20 cells were quantified across 3 biological replicates per condition. Each point on the graph represents a condition replicate. Error bars represent mean +/- SEM. Statistics: Unpaired two-tailed t-test. *p<0.05, **p<0.01. **C)** DGAT mechanism of action. **D)** IMR90s were treated with DGAT1/2 inhibitor for 24 hours, followed by 6 hours of 10 µM etoposide treatment. Statistics: one-way ANOVA with Tukey’s post hoc test. Error bars on graphs indicate mean +/- SEM. ***p<0.0005, ****p<0.0001. **E)** Senescence marker readouts in IMR90s pre-treated with DGAT inhibitors. IMR90s were treated with 10 µM etoposide for 48 hours, followed by 5 days of recovery. Senescence-associated beta-galactosidase staining (left) and RT-qPCR for cell cycle arrest markers (right). Cells were kept in the DGAT1/2 inhibitor for the duration of the senescence induction. Statistics: one-way ANOVA (comparisons within genes). Error bars on graphs indicate mean +/- SEM. **p<0.01, ***p≤0.001, ****p<0.0001.

Given the pro-inflammatory roles for LDs and the benefits of their ablation in inflammatory contexts ^19–21^, we tested whether ablating LDs before genotoxic stress would affect the response to DNA damage, hypothesizing this would reduce DNA damage burden and markers of senescence. We tested this by blocking triglyceride (TAG) formation by inhibiting the DGAT enzymes, which are responsible for TAG synthesis from diacylglycerol (DAG) (**Fig 1C**). As the knockout of DGAT1 or DGAT2 alone still allows LDs to form, we chose to inhibit both DGAT enzymes to completely block LD biogenesis. DGAT1/2 inhibition for 24 hours was sufficient to starkly reduce LDs in both low-passage (P10; low endogenous LD levels) and high-passage (P25; LD-rich) IMR90 fibroblasts, at doses that do not induce DNA damage (γH2Ax) (**Supp Fig 1A, 1B**). We then treated IMR90s with DGAT1/2 inhibitors for 24 hours, followed by 6 hours of high-dose etoposide (25 µM). Contrary to our hypothesis that reduced LDs and subsequently reduced inflammation would attenuate cellular senescence, we initially observed no effect on senescent cell accumulation (**Supp Fig 1C**). We then reasoned that high doses of etoposide in IMR90s could potentially overwhelm the cells’ DNA repair capacity, and thus trialed lower etoposide doses. We observed a notable increase in senescence, as measured by SA-β- galactosidase and *CDKN1a/CDKN2a* expression, at lower etoposide doses relative to higher doses, suggesting a DSB range of pharmacologic intervention *in vitro*, beyond which the DNA damage burden is too high (**Supp Fig 1D,E**). IMR90 cells pre-treated with DGAT1 and DGAT2 inhibitors showed more DNA damage than cells treated with etoposide alone (**Fig 1D**). Further, cells pre-treated with DGAT1/2 inhibitors showed increased senescence markers, as measured by SA-β-galactosidase staining and by the expression of cell cycle arrest genes, *CDKN1A* (p21) and *CDKN2A* (p16) (**Fig 1E**), suggesting that LD ablation worsens the long-term consequences of acute DNA damage.

### ATGL overexpression reduces DNA damage *in vitro* and *in vivo*

Evidence in the literature suggests that cells upregulate fatty acid oxidation and oxidative phosphorylation after DNA damage, while deprioritizing glycolysis ^22,23^. One possible explanation for the effect of DGAT1/2 inhibition on DNA damage is that, by preventing LD accumulation, cells are deprived of substrates for lipolysis and fatty acid oxidation, potentially interfering with the DNA damage response. To test whether enhanced lipolysis confers a benefit against DNA damage, we overexpressed ATGL to promote lipolysis in cells prior to DNA damage. Remarkably, cells overexpressing ATGL showed reduced DNA damage at multiple time points of etoposide treatment (**Figs 2A,B**). While etoposide induces DSBs in all stages of the cell cycle, it preferentially causes DNA damage during the S/G2 phases, resulting in G2/M stalling ^24,25^. Though we observed no effects of ATGL overexpression on the cell cycle (**Supp Fig 2A**), we sought to test if ATGL overexpression still conveyed benefits with an insult that causes DNA damage agnostic of the cell cycle. Thus, we used gamma irradiation, which causes robust DSBs independent of the cell cycle. As expected, 10 Gy of gamma irradiation robustly increased DNA damage in both control and ATGL overexpressing cells immediately after irradiation (**Figs 2C,D**). However, after 24 hours of recovery, DNA damage in ATGL-overexpressing cells returned to unirradiated levels, whereas those expressing the GFP control showed limited recovery (**Fig 2D**). We also observed this DNA damage reduction in primary MEFs treated with etoposide, suggesting this effect is not cell type-specific; primary MEFs overexpressing ATGL show a reduction in total γH2Ax-positive cells, as well as cells with a high burden of DNA damage (**Fig 2E**). Of note, total histone H2Ax is unchanged with ATGL overexpression, suggesting differences in γH2Ax are a result of changes in DNA damage and not variations in histone abundance (**Supp Fig 2B**).

**Figure 2.**
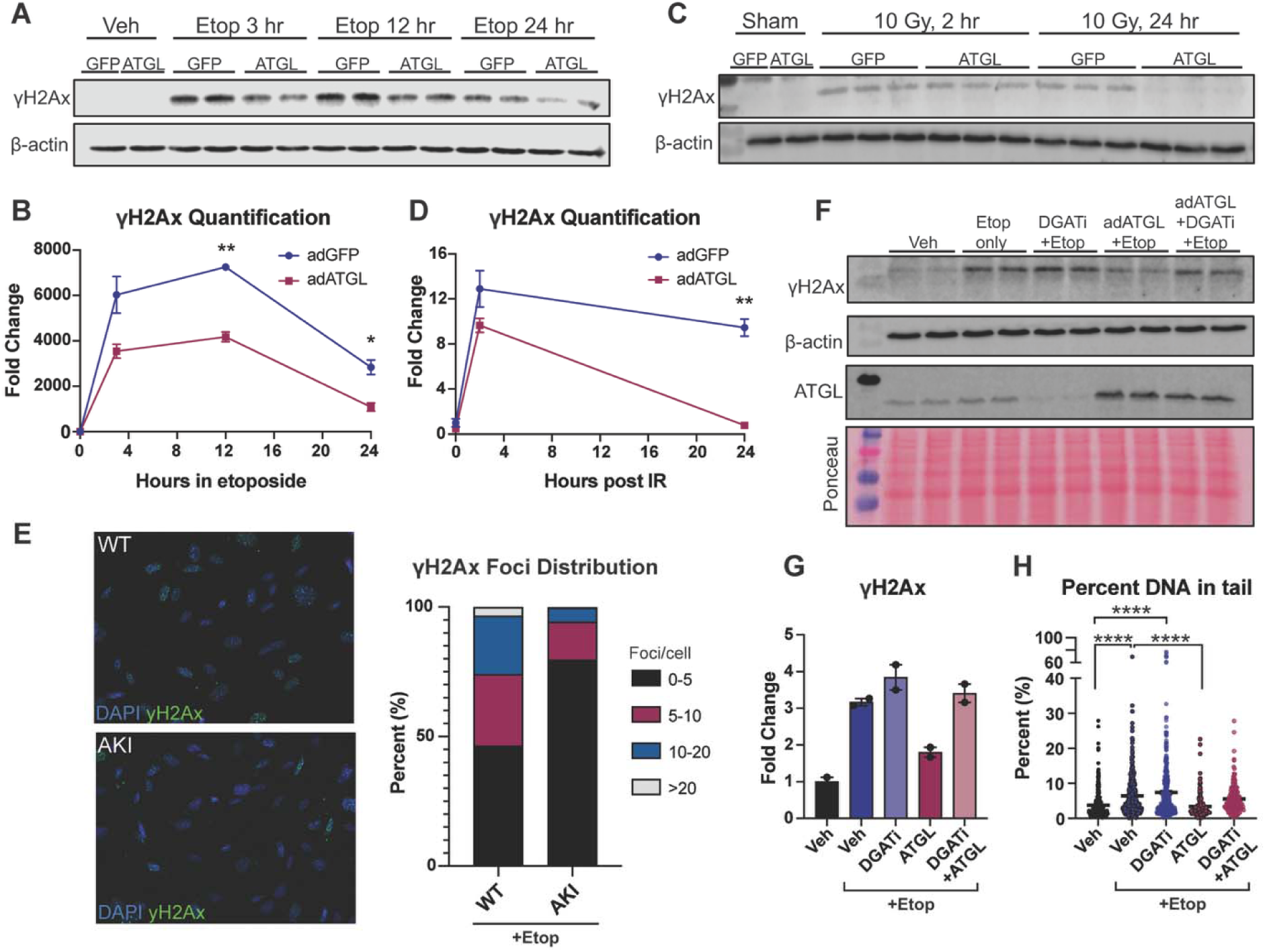
ATGL overexpression reduces DNA damage. **A)** AML12 cells transduced with ATGL or GFP control adenoviruses were treated with 30 µM etoposide for the indicated times and assayed for γH2Ax. **B)** Quantification of A. *p<0.05, **p<0.01. **C)** AML12 cells transduced with ATGL or GFP control adenoviruses were irradiated (10 Gy) and harvested at the indicated times post-IR. **D)** Quantification of C. **p<0.01. **E)** Primary MEFs were isolated from WT or ATGL knock-in (AKI) mice, followed by etoposide incubation for 6 hours. Immunofluorescence was performed for γH2Ax, and cells were imaged using the Agilent Gen5 Cytation. Representative γH2Ax images and foci distribution are shown. **F)** Cells were treated with DGAT inhibitors for 24 hours, followed by ATGL adenoviral transduction, followed by 24 hours of etoposide treatment. Western blotting for γH2Ax, β-actin, and ATGL. **G)** Quantification of F. **H)** AML12 cells were treated as in F. The Comet assay was performed on cells after 24 hours of etoposide. Comets were quantified with OpenComet software. A minimum of 180 comets were quantified per condition. Statistics: one-way ANOVA with Tukey’s post hoc test. ****p<0.0001.

To confirm that this effect on DNA damage was not an off-target effect of ATGL overexpression, we pretreated AML12 cells with DGAT inhibitors to ablate LD formation, then overexpressed ATGL. Importantly, overexpression of ATGL reduced γH2AX in etoposide-treated cells, but this effect was abrogated by pretreatment with DGAT1/2 inhibition (**Fig 2F,G**). Further, we performed neutral comet assays to directly assess DNA breaks. In agreement with γH2Ax data, ATGL overexpression reduced DNA damage as measured by neutral comet assay (% DNA in tail), an effect abrogated by pretreatment with DGAT inhibitors (**Fig 2H**). Together, these data show that the presence of LDs is required for the benefit conferred by ATGL overexpression, suggesting that enhanced LD breakdown (lipolysis) underlies the benefit of ATGL-mediated DNA repair.

To test these effects *in vivo*, we generated mice globally overexpressing ATGL. A construct harboring pAAV-CMV-Beta actin-LOX-STOP-LOX-ATGL FLAG was generated and inserted in the Rosa locus of C57BL/6 mice using a TILD-CRISPR strategy. These mice were crossed with mice expressing E2A Cre (embryonic Cre activation) to generate global ATGL-overexpressing mice carrying a single additional copy of ATGL (“AKI”) (**Fig 3A,B**). While livers from AKI mice are histologically similar to WT controls, they have a reduced LD burden, as measured by Oil Red O staining, indicating enhanced lipolysis in these mice (**Fig 3C,D**). Further, Primary MEFs harvested from AKI mice show resistance to LD accumulation upon oleic acid loading (**Fig 3E**), indicating enhanced lipid turnover expected in cells with high lipolytic activity. We performed dose-response and time-course analyses of gamma irradiation in WT mice and determined that 5-10 Gy of irradiation was sufficient to induce DNA damage in multiple tissues and that the γH2Ax signal was substantially reduced by 6 hours post-irradiation (**Supp Fig 2C,D**). To assess differences in DNA damage prior to full recovery, we irradiated mice with 7.5 Gy and allowed for 4 hours of recovery. Consistent with cell culture data, AKI mice showed less γH2Ax at 4 hours post-irradiation in all measured tissues, including the liver, lung, kidney, and heart, relative to Cre-negative sham controls (**Fig 3F**), suggesting that the effects of ATGL are conserved across cell types.

**Figure 3.**
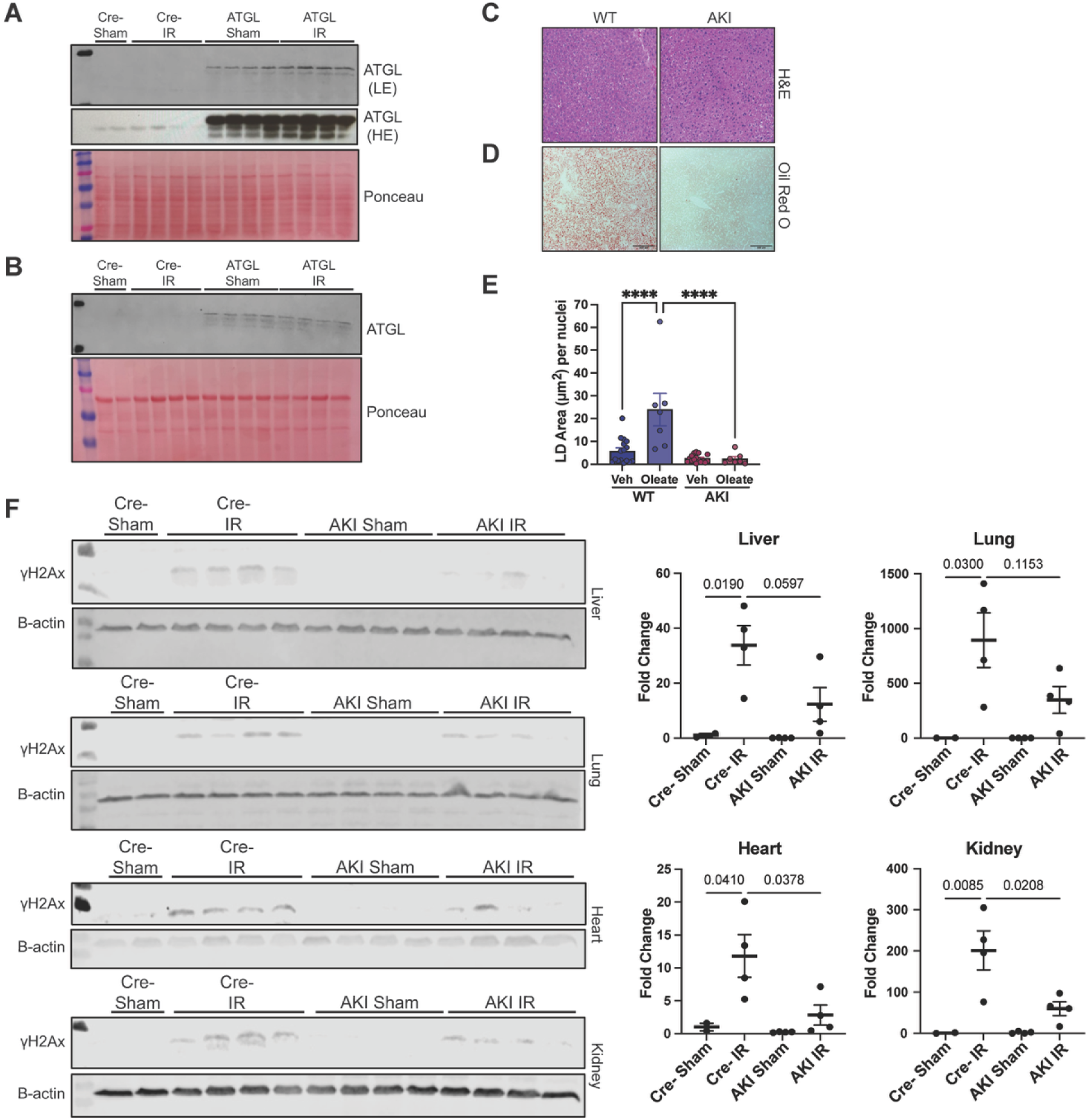
ATGL overexpression reduces DNA damage in vivo. A, **B)** Validation of ATGL overexpression in E2A Cre-AKI liver (A) and lung (B). **C**) H&E staining of WT and AKI livers. **D)** Oil Red O staining of WT and AKI livers. Scale bar = 100 micron. **E)** Primary MEFs were treated with 100 µM oleate overnight to induce LD accumulation, followed by one hour of oleate washout. Cells were stained with BODIPY 493/503 and quantified. **F)** γH2Ax results and quantification from tissues harvested from E2A Cre-ATGL (AKI) mice irradiated with 7.5 Gy + 4- hour recovery. Mice were of mixed sexes, n=4 mice. P-values from one-way ANOVA are displayed.

### ATGL mitigates the downstream consequences of DNA damage

Since ATGL reduces acute DNA damage, we next asked if this protection translates to beneficial long-term outcomes. Our lab previously showed that global overexpression of the ATGL homolog *bmm* in *Drosophila melanogaster* improved multiple measures of physiological fitness and reduced markers of aging, including locomotion, fecundity, and stress resistance ^26^. As senescence is a hallmark of aging and ATGL reduces DNA damage (a senescence precursor), we hypothesized that ATGL overexpression would protect against entry into senescence. We repeatedly passaged WT and AKI Primary MEFs to induce replicative stress. At passage 6, WT MEFs had accumulated approximately 75% β-gal positive cells, while AKI MEFs of the same passage number accumulated less than half the number of senescent cells (**Fig 4A**). Similarly, AKI MEFs were refractory to the increase in cell cycle arrest and senescence-associated secretory phenotype (SASP) gene expression downstream of etoposide treatment (**Fig 4B**). Together, these data suggest ATGL confers resistance to a senescent state in primary MEFs.

**Figure 4.**
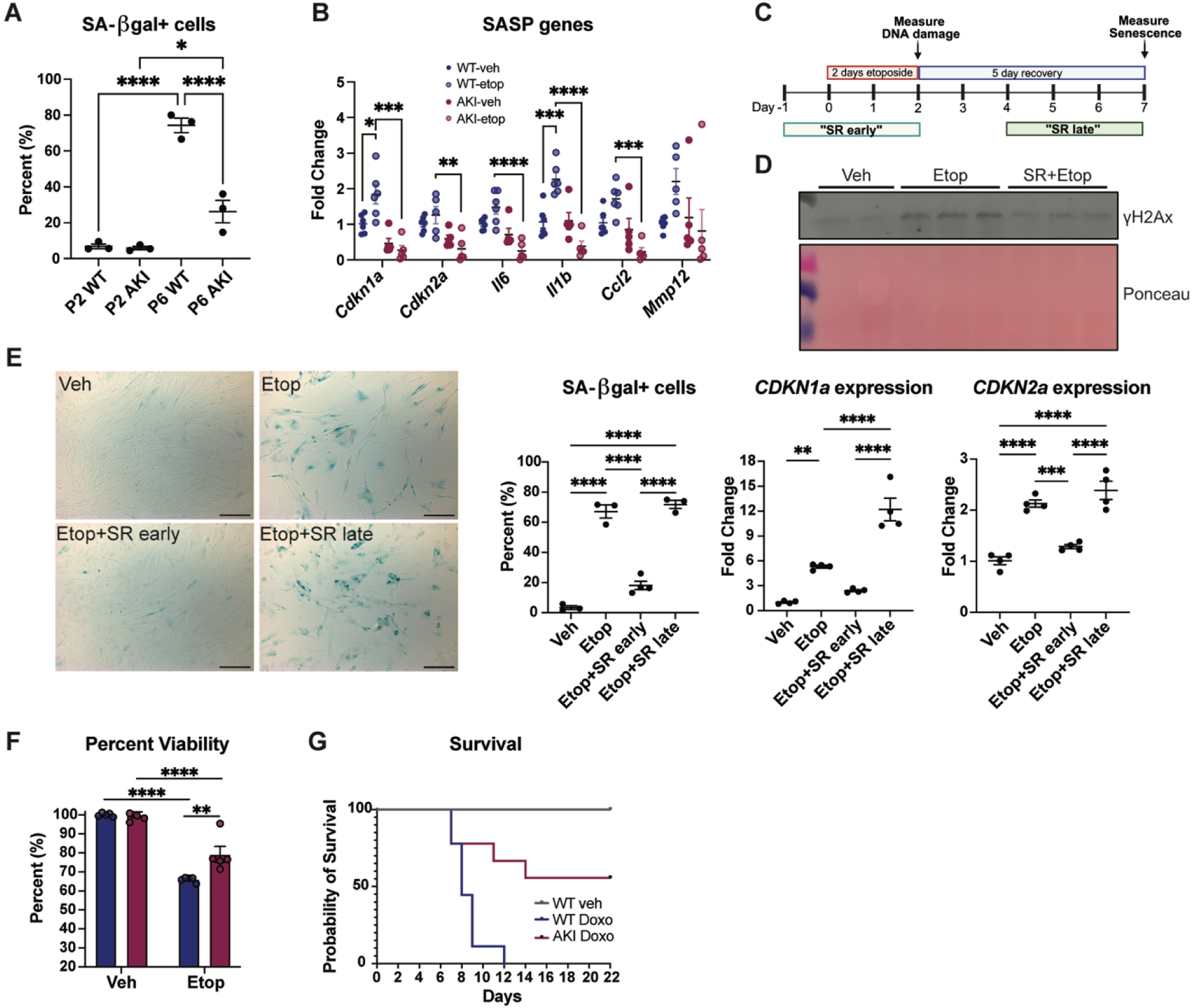
ATGL/lipolysis reduces markers of cellular senescence. **A)** Senescence-associated β-galactosidase measurement of high-passage primary MEFs isolated from WT or AKI mice. Statistics: one-way ANOVA. *=p<0.05, ****=p<0.0001. **B)** AKI and WT primary MEFs were treated with 10 µM etoposide for 24 hours, followed by 5 days of recovery to induce senescence. Senescence-associated secretory phenotype (SASP) gene expression measured by RT-qPCR. **C)** Schematic of SR4995 timing experiment in IMR90s. **D)** γH2Ax measurement after 2 days of etoposide treatment. **E)** Senescence measurements denoted at the timepoint shown in (C). Scale bar = 200 microns. **F)** Viability measurement of WT and AKI primary MEFs treated with 30 µM etoposide for 60 hours using the Alamar Blue assay. Statistics: one-way ANOVA with Tukey’s post hoc test. **=p<0.01, ****=p<0.0001. **G)** Survival after 20 mg/kg IP injection of doxorubicin hydrochloride in WT and AKI mice. n=6-9.

Our models of ATGL overexpression involve either genetic overexpression or adenoviral transduction, both of which induce ATGL expression before DNA damage. Therefore, we sought to test whether this protective effect on genomic stability is specific to the timing of enhanced lipolysis. To do this, we utilized SR4995, a known lipolysis activator. In a basal state, the ATGL coactivator protein CGI-58 (alias ABHD5) is bound to a perilipin protein, thereby preventing ATGL activation ^27^. SR4995 binds CGI-58, promoting its dissociation from perilipin proteins and enabling binding of ATGL and subsequent activation of ATGL to promote lipolysis ^28^. Given the robust effects of SR4995 on DNA damage mitigation (**Supp Fig 3C,D**), we utilized this drug to modulate lipolysis in a time-dependent manner. We treated IMR90 cells with SR4995 before DNA damage (“SR early”) or after DNA damage (“SR late”) (**Fig 4C**). Consistent with previous experiments, SR4995-pretreated IMR90s showed reduced γH2Ax after a two-day etoposide incubation (**Fig 4D**). Interestingly, we observed a reduction in β-gal positive cells and cell cycle arrest markers (markers of cellular senescence) with SR4995 treatment, but only when SR4995 was administered before DNA damage; enhancing lipolysis after DNA damage did not confer a benefit (**Fig 4E**). AKI MEFs also demonstrated higher 60-hour cell viability in response to high-dose etoposide, suggesting resistance to etoposide-induced cell death (**Fig 4F**). Given the genoprotective effect of ATGL, we next tested the robustness of these effects by administering a lethal dose of doxorubicin hydrochloride, a commonly used antineoplastic drug and a source of acute DNA damage in humans undergoing chemotherapy, to both control and ATGL transgenic mice. While all wild-type mice were deceased within 12 days after administration of doxorubicin, more than half of the AKI mice survived this systemic genotoxic challenge (**Fig 4G**). Together, these data show that enhanced lipolysis primes tissues to respond more effectively to acute genotoxic stress, thereby preventing senescence and enhancing survival.

### ATGL and lipolysis promote DSB repair

Given that ATGL promotes γH2Ax resolution and minimizes long term consequences of DNA damage, we sought to further investigate ATGL’s role in DNA DSB repair by measuring markers of HR and NHEJ. To assess HR, we measured nuclear Rad51 foci 24 hours after etoposide removal, which indicate Rad51 localization to chromatin and the progression of homology-directed repair. ATGL-overexpressing cells had increased total Rad51 foci, with a trending decrease in total γH2Ax foci (**Fig 5A-C**). Further, ATGL increased the overlap of Rad51 and γH2Ax signal, indicating that the sites of damage had recruited more Rad51 to facilitate repair (**Fig 5D**). To interrogate whether ATGL promotes NHEJ, we performed chromatin enrichments to assess how ATGL changes the localization of the NHEJ-promoting proteins. Interestingly, ATGL promotes 53BP1 recruitment to the chromatin acutely after DNA damage (3 hours post-etoposide) (**Fig 5E**). ATGL also increases chromatin-bound 53BP1 even without DNA damage, suggesting a priming effect, in which lipolysis activation prior to damage gives the cell a greater capacity to repair the DNA once damage occurs (**Fig 5E**). We corroborated this baseline induction of 53BP1 using a 53BP1 reporter (**Supp Fig 3A**). To functionally test whether NHEJ-mediated repair was occurring, we employed the widely used NHEJ reporter assay, which uses a fluorescent construct (pimEJ5GFP) containing an Sce-I cut site. When transfected into cells, the plasmid encoding the Sce-I endonuclease cuts at its designated site, resulting in GFP fluorescence upon successful in-frame NHEJ repair. When measured by flow cytometry 48 hours after transfection with the Sce-I plasmid, we observed a marked increase in GFP-positive cells when cells were pretreated with SR4995 (**Fig 5F**). We were able to demonstrate this induction in NHEJ in both liver cells and human fibroblasts, suggesting this effect is conserved across cell types (**Fig 5G**). Further, pre-treatment with a DNA-PK inhibitor (inhibits NHEJ) and Rad51 inhibitor (inhibits HR) abrogated the benefit of ATGL on γH2Ax burden at doses confirmed not to induce DNA damage (**Fig 5H, Supp Fig 3B**). Together, these data suggest that ATGL mitigates DNA damage by enhancing DNA repair mechanisms.

**Figure 5.**
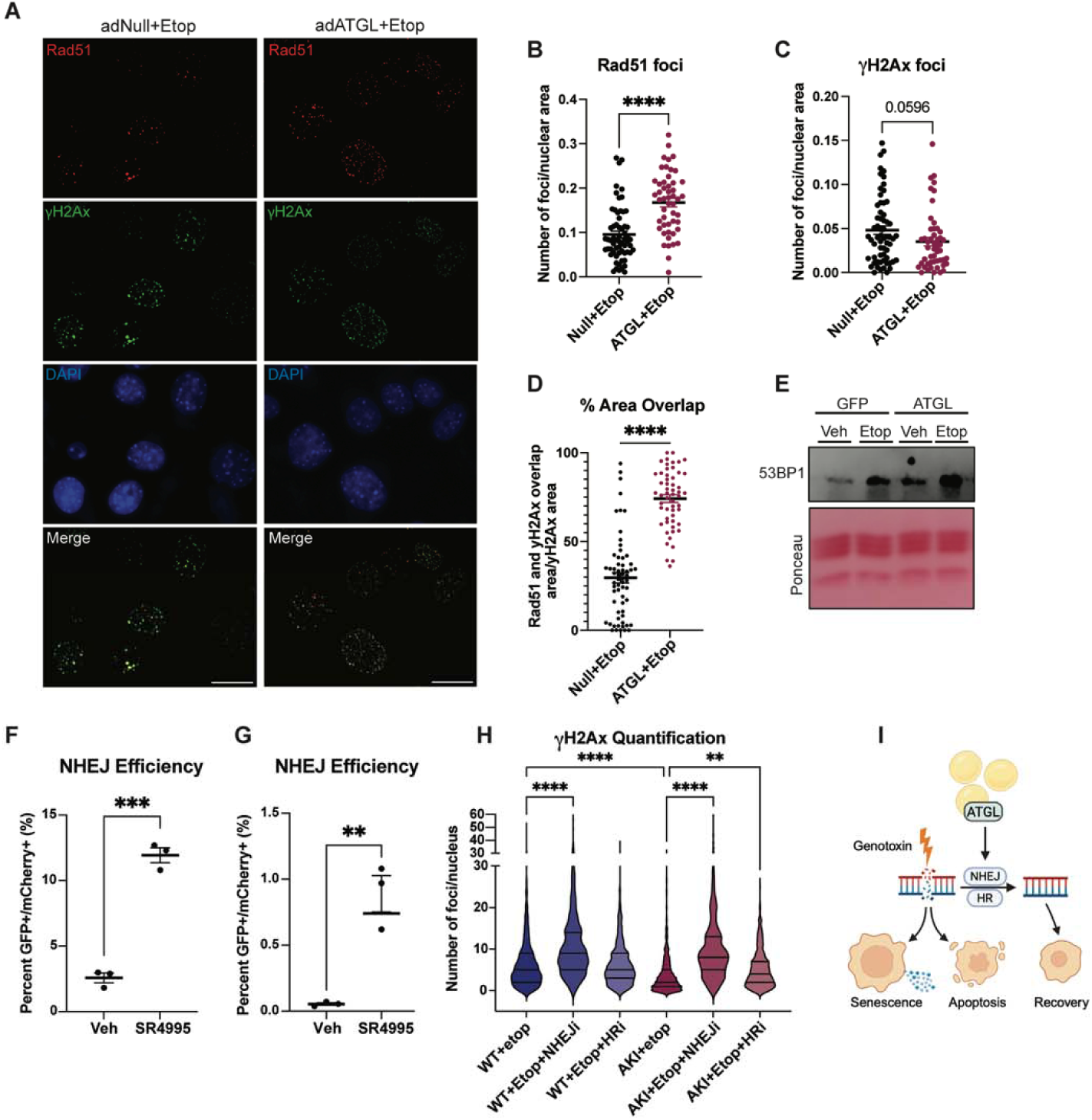
ATGL/lipolysis promote DNA repair. **A)** AML12 cells were transduced with adNull or adATGL, treated with etoposide for 3 hours, and assayed for γH2Ax and Rad51 immunofluorescence after 24 hours of recovery. Representative images and foci number quantification per nucleus. Scale bar = 25 microns. **B)** Quantification of Rad51 foci. n=70-80. Each point on the graph represents a single nucleus. Statistics: Unpaired t-test. ****=p<0.0001. **C)** Quantification of γH2Ax foci. Same nuclei as quantified in B (n=70-80). Statistics: Unpaired t-test. **D)** Quantification of γH2Ax/Rad51 area overlap. The percent area overlap of the cells quantified in B and C was calculated using the BIOP JACoP macro in FIJI. Statistics: Unpaired t-test. ****=p<0.0001. **E)** 53BP1 immunoblot on chromatin fractions. Cells were treated with etoposide for 3 hours, then harvested and enriched for chromatin. Representative western blot of 2 independent experiments. **F)** Assessment of NHEJ repair using pimEJ5-GFP reporter that was stably transfected into AML12 cells. Statistics: one-way ANOVA with Tukey’s post hoc test. Error bars on graphs indicate mean +/- SEM. **G)** Assessment of NHEJ repair using HCA2-NHEJ I9a human fibroblast cell line. Statistics: Unpaired t-test. **=p<0.005. **H)** Primary MEFs were pretreated with an NHEJ inhibitor (Nu7441 at 10 µM) or an HR inhibitor (RI-1 at 20 µM) for 1 hour, followed by 3 hours of 10 µM etoposide and 6 hours of recovery. Cells were then fixed and stained for γH2Ax, followed by foci quantification. Statistics: One-way ANOVA. **=p<0.01, ****=p<0.0001.

## DISCUSSION

Here, we demonstrate that ATGL modulates the cellular response to acute DNA damage, bridging two seemingly distinct biological processes: lipid metabolism and DNA repair. By demonstrating this link, our findings implicate ATGL and lipid catabolism as upstream regulators of the pathways governing genomic integrity and cell survival.

While the intersection of lipid metabolism and DNA repair remains relatively understudied, supporting evidence is emerging. Existing data indicate that cells undergo an acute metabolic shift away from glycolysis and toward fatty acid oxidation and oxidative phosphorylation immediately following DNA damage ^23,29,30^, Consistently, inhibiting fatty acid oxidation exacerbates both acute DNA damage and its long-term consequences, such as cellular senescence, underscoring the importance of this metabolic pivot in mitigating genotoxic stress ^31^. Further, lipid metabolism has been shown to support DNA repair by promoting the acetylation of key repair proteins, providing a direct biochemical link between lipid catabolism and genomic stability ^30^.

Although we investigated ATGL primarily through the lens of acute DNA damage, unrepaired lesions inevitably promote chronic genomic instability - a primary driver of senescence, mutagenesis, and age-related disease. Because our data show that ATGL-mediated lipolysis reduces senescence, this work carries broad physiological implications beyond acute, exogenous genotoxic stress. Fasting and caloric restriction, which physiologically upregulate ATGL and lipolysis, are potent enhancers of healthspan and longevity. Our findings suggest that downstream effects of fasting, specifically lipolysis, essentially “prime” cells to mitigate DNA damage, offering a novel mechanism for how dietary restriction preserves genomic stability.

Fasting is well documented to protect mice against lethal levels of DNA damage ^32–34^. Emerging evidence highlights that fasting exerts a dual effect: it is cytoprotective for normal cells but cytotoxic to transformed cells. For instance, fasted mice bearing pancreatic tumors subjected to irradiation exhibit dramatically prolonged survival and delayed tumor progression compared to their fed counterparts ^34^. Similarly, dietary restriction selectively sensitizes cancer cells to chemotherapy while shielding healthy tissues ^35^. Given that ATGL is required to mediate the longevity benefits of dietary restriction ^36^, our work suggests that lipolytic activity is a key component of the beneficial responses to fasting during genotoxic stress.

Our findings are highly consistent with the known metabolic shifts that accompany aging. Aging is characterized by a systemic decline in lipid catabolic pathways ^37–40^, leading to ectopic lipid droplet accumulation (reviewed in ^41^). Conversely, elevated fatty acid oxidation is the most prominent metabolic signature of centenarians ^42^, and robust mitochondrial β-oxidation best distinguishes individuals experiencing healthy aging from rapid agers ^43^. Synthesizing these observations with our current data, it is plausible that the age-dependent decline in lipolysis compromises genomic stability, thereby accelerating the aging process. This is further supported by our finding that overexpressing the ATGL homolog in *C. elegans* promotes multiple indices of healthspan ^26^, reinforcing a conserved, beneficial role for lipid catabolism in longevity.

Importantly, cell-intrinsic lipolysis positively regulates several other hallmarks of aging, including macroautophagy ^44,45^, nutrient sensing ^13,46^, epigenetics ^13,47,48^, mitochondrial function ^13,49,50^, and intercellular communication ^51,52^. Consequently, ATGL may influence DNA repair through a network of interconnected downstream pathways. Future studies will be required to dissect these specific mechanisms and determine ATGL’s contribution to various forms of DNA damage and repair pathways beyond double-strand breaks.

In summary, we identify an important role for ATGL and pharmacological lipolysis activation in promoting DNA repair, enhancing cell survival, and preventing senescence in response to genotoxic stress. These findings emphasize the contribution of lipid metabolism to genomic preservation and uncover new therapeutic avenues for protecting genomic stability.

## ACKNOWLEDGEMENTS

We would like to thank the University of Minnesota Imaging Centers and the Masonic Institute for the Biology of Aging and Metabolism for providing instrumentation and expertise. Grants that supported this work include NIH grants F30AG076322 (MD), R01HL163011 (HDN), R01DK132849 (DGM), R01AG069768 (DGM), T32AG029796 (MD, RKM), T32DK007203 (EK), Masonic Cancer Center (HDN), Edward P. Evans Foundation (HDN), American Society of Hematology (HDN), AACR Career Development Award (HDN).

## METHODS

### Mice

All animal protocols were approved by the University of Minnesota Institutional Animal Care and Use Committee. All mice were maintained on a chow diet (2018 Teklad Global 18% Protein Rodent Diet) with access to water *ad libitum* under a 14/10 light/dark cycle and controlled temperature (23L±L1°C). WT and ATGL floxed mice were of C57/BL/6J background. E2A Cre mice (B6.FVB-Tg(EIIa-Cre)C5379Lmgd/J, Jackson Labs) were of B6.FBV background. For the generation of ATGL floxed mice, a construct harboring pAAV-CMV-Beta actin-LOX-STOP-LOX-ATGL FLAG was generated and inserted into the Rosa locus of C57BL/6 mice using a TILD-CRISPR strategy. These mice were crossed with mice expressing E2A Cre (embryonic Cre activation) to generate global ATGL-overexpressing mice carrying a single additional copy of ATGL (AKI).

### Irradiation

Irradiation (IR) experiments were performed using an RS2000 X-ray irradiator. Dose-response and time course studies were performed to determine the appropriate dose of IR to induce a γH2Ax response in tissues of interest. Mice underwent total body irradiation at 7.5 Gy, single exposure, with mouse harvest 4 hours post-IR. Mice did not receive specialized food or prophylactic antibiotics given the short time frame between irradiation and harvest. Body weights for all mice were normalized across groups for IR studies so that similar absorbed radiation doses were obtained. Sham control mice were placed in the irradiator for the same duration as irradiated mice, without turning on the irradiator.

### Cell culture

All cells, except primary mouse embryonic fibroblasts (MEFs), were maintained in a humidified incubator at 37°C, 5% CO_2_. AML12 cells obtained from ATCC were cultured in DMEM:F12 media supplemented with 10% FBS, 1% ITS (10 mg/mL insulin, 5.5 mg/mL transferrin, 5 ng/mL selenium), 1% Pen/strep, and 50 ng/mL dexamethasone. IMR90 cells were cultured in MEM, 10% FBS, 1% Pen/strep, and 1% nonessential amino acids. IMR90s were only used up to passage 18 for non-replicative senescence experiments. Primary MEFs were isolated as described.^53^ Briefly, breeding pairs were mated and monitored for a vaginal plug. At E13.5 (13 days after plug), pregnant females were euthanized by CO_2,_ and the pregnant uterus was quickly dissected. Embryos were decapitated to minimize the contribution of neural tissue to MEF population, and the liver and heart were dissected and removed. The remaining embryonic tissue was manually minced, digested in 0.25% trypsin, and subsequently plated in culture media. Primary MEFs were cultured in 50% DMEM/50% F10 media, supplemented with 10% FBS, 1% nonessential amino acids, 1% pen/strep, and 1% Glutamax. Cells were cultured in a humidified incubator at 37°C in 3% O_2_/5% CO_2_ for expansion. Cells were brought to standard culture conditions (5% CO_2_, atmospheric oxygen) 24 hours before initiating experiments. Primary MEFs were only used up to passage 4 for non-replicative senescence experiments.

For adenoviral transductions, ATGL (Lot: 20160819, mouse ATGL) was obtained from VectorBiolabs, TrackGFP (adGFP) was obtained from UNC Gene Therapy Center, and Null adenovirus was obtained from VectorBiolabs (#1300). AdGFP was used for the control for all non-imaging experiments, and adNull was used for Rad51/p53 imaging experiments and Cytation experiments. Adenoviral transductions were performed in serum-free media to an equivalent multiplicity of infection for 6 hours, at which point media containing FBS was added to each well. The next morning, the viral media was replaced with fresh media containing FBS. Etoposide treatments were performed 48 hours after the start of transduction.

### Chemical reagents

A list of chemical reagents can be found in Table 1.

**Table 1.** Chemical Reagents.

| Reagent | Dose |
| --- | --- |
| Etoposide (Millipore 341205) | 30 µM for acute DNA damage. 10 µM for senescence induction |
| SR4995 (Sigma SML2207) | 5 µM |
| ATMi (KU55933) | 10 µM |
| ATRi (AZD6738) | 2 µM |
| Nu7441 (Tocris 3712) | 10 µM |
| RI-1 (SelleckChem S8077) | 20 µM |

### Real-time quantitative PCR

Cells were lysed with Trizol reagent and RNA was extracted using phenol/chloroform separation. RNA was reverse transcribed to cDNA with the qScript cDNA Synthesis Kit (Quantbio), and then cDNA was amplified using SYBR Green Master Mix for qPCR (Applied Biosystems). β-actin was used as a housekeeping gene. A list of primers can be found in Table 2.

**Table 2.**
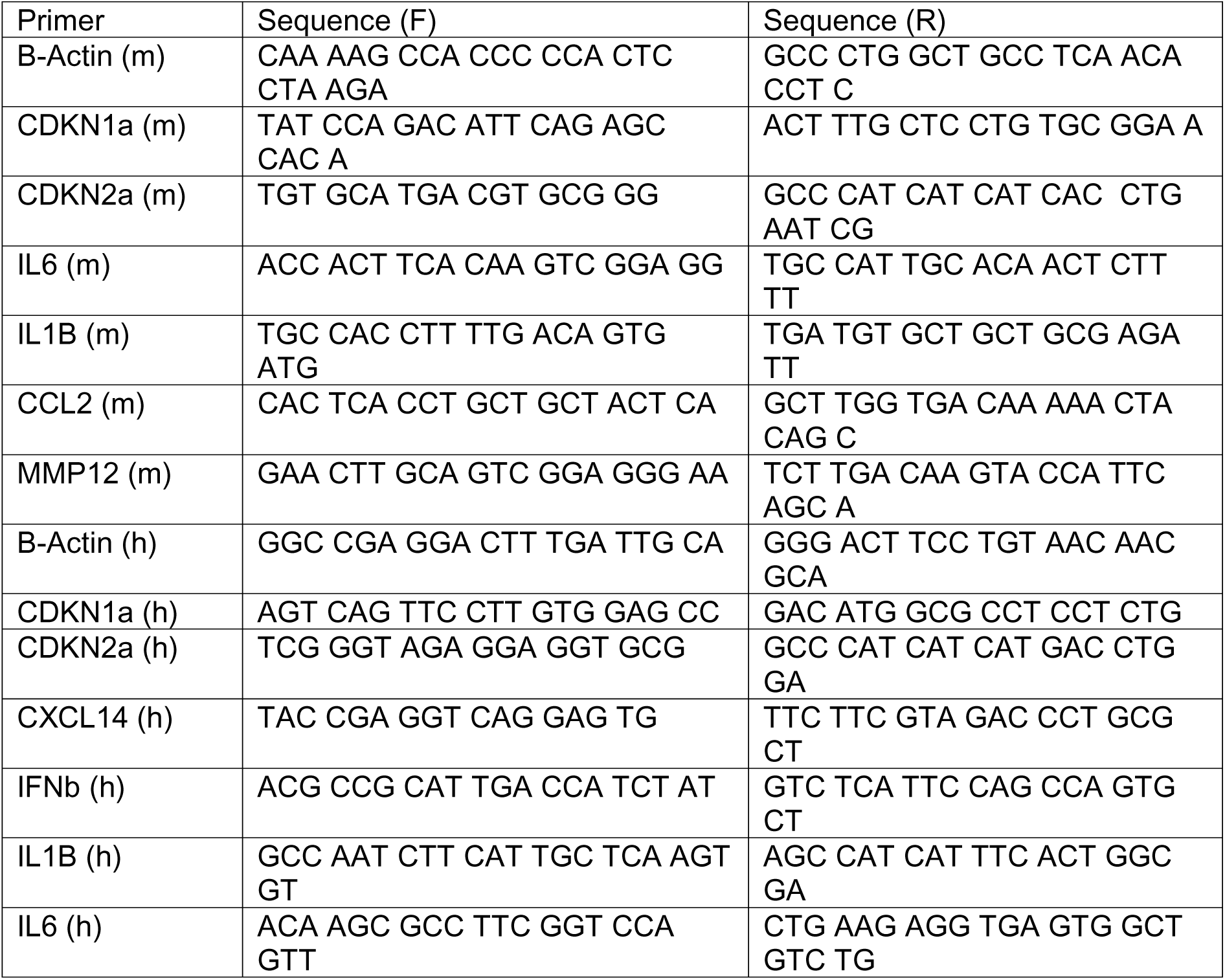
RT-qPCR Primers.

### NHEJ reporter assay

The pimEJ5-GFP plasmid from Addgene (44026) was transfected into AML12 cells and selected using puromycin for 2 weeks. Cells were then transduced with adATGL or adNull virus as a control. The following day, cells were transfected with the I-SceI plasmid and mCherry control. Cells were analyzed by flow cytometry 48 hours later. For SR4995 conditions, cells were treated with SR4995 starting the day of Sce-I transfection and maintained in SR4995 for the duration of the experiment. The NHEJ efficiency was calculated as the ratio between GFP+ cells/mCherry+ cells. The same experiments were performed with the NHEJ I9a reporter in HCA2 fibroblasts, which were obtained from Vera Grobunova ^10^.

### Immunofluorescence and Imaging

Cells were grown on collagen-coated glass coverslips. Cells were treated with relevant drug pre-treatments, followed by 30 µM etoposide for 3 hours, then recovered in fresh media for 24 hours prior to staining. For immunostaining Rad51/γH2Ax, samples underwent brief pre-extraction, using 0.1% Triton-X for 2 minutes at 4°C. Samples were immediately fixed with 4% PFA for 15 minutes, and permeabilized with 0.5% Triton-X for 15 minutes at room temperature. Samples were blocked with 5% BSA or goat serum for 1 hour, followed by overnight antibody incubations. Antibody concentrations were: Rad51 (ab133534), 1:200; γH2Ax (Cell Signaling 9719S), 1:250. Images were taken on a Leica DMi8 epifluorescence scope. For foci quantification, nuclear ROIs were defined, and foci within nuclear ROIs were identified using the “find maxima” function in FIJI. The percent area overlap of Rad51 and γH2Ax signal was calculated using the BIOP JACoP macro in FIJI.

For 53BP1 imaging, AML12 cells were stably transfected with mApple-53BP1 (Addgene #69531). Images were acquired on the Lionheart FX with a 20x objective. WT AML12 cells were used as controls. For LD staining, cells were fixed with 4% PFA for 5 minutes, followed by 3 washes in PBS. BODIPY 493/503 staining was then performed for 20 minutes at 2 µM.

### Senescence staining

The Senescence-Associated β-Gal Staining Kit was purchased from Millipore (CS0030) and was performed according to the manufacturer’s instructions. Cells were treated with 20 µM etoposide for 2 days and allowed to recover for 5 days. Cells were fixed using the kit’s fixative and stained with Xgal solution for 12 hours.

### Western blotting

Cells were harvested in lysis buffer (150 mM NaCl, 10 mM Tris-HCl, and 1% Triton X). Before protein determination, lysates were sonicated 2x at 15% amplitude at 4°C. Protein lysates were subjected to SDS-polyacrylamide gel electrophoresis and transferred to PVDF membrane before immunoblotting. Membranes were incubated with primary antibody overnight at 4°C. Antibodies used were: γH2Ax (Bethyl A300-081A) at 1:2000; ATGL (Cell Signaling 2138S) at 1:1000, β-actin (A2228) at 1:5000, 53BP1 (Ab36823) at 1:1000, pan-acetylated lysine (Cell Signaling 9681S) at 1:1000.

### Comet assay

Neutral comet assays were performed on AML12 cells using R&D Biosystems 4250-050-K kit. Cells were transduced with an adATGL adenovirus or an adNull control. Forty-eight hours after transduction, cells were treated with 30 µM etoposide. After 24 hours of etoposide treatment, the comet assay was performed following the manufacturer’s instructions. Images were analyzed using OpenComet software.

### Cell Viability

Cells were seeded in a 96-well plate. After drug treatments, wells were incubated in 30 µM etoposide for 60 hours. Cell Viability was determined using the Alamar Blue assay (Invitrogen DAL1025). Fluorescence intensity at 580 nM was measured 2-4 hours after the Alamar Blue reagent was added to cells.

### Chromatin Fractionation

A method for chromatin fractionation was largely adapted from a previously published protocol^54^. Briefly, cells were lysed with 5 volumes of E1 buffer (50 mM HEPES-KOH, pH 7.5, 140 mM NaCl, 1 mM EDTA, pH 8.0, 10% glycerol, 0.5% NP-40, 0.25% Triton X-100, 1 mM DTT), complemented with 1x protease inhibitor, phosphatase inhibitor, and deacetylase inhibitor cocktail. The supernatant was collected as the cytoplasm fraction, and the pellet was washed with E1 buffer twice with centrifugation in between. The pellet was resuspended by gentle pipetting in 2 volumes of ice-cold E2 buffer (10 mM Tris-HCl, pH 8.0, 200 mM NaCl,1 mM EDTA, pH 8.0, 0.5 mM EGTA, pH 8.0), complemented with 1x protease and phosphatase inhibitor, and deacetylase inhibitor, and spun at 1100xg at 4°C for 2 min. The pellet was resuspended in the same volume of E2, an aliquot was taken as the nuclear fraction, and centrifuged as before. The pellet was washed in E2 buffer twice more with centrifugation at 1100xg for 2 minutes in between. The supernatant was discarded, and the pellet was resuspended in ice-cold E3 buffer (500 mM Tris-HCl, 500 mM NaCl), complemented with 1x protease inhibitor cocktail, phosphatase inhibitor, and deacetylase inhibitor. The solution (chromatin fraction) was sonicated 3X at 20% amplitude for 10 seconds. Fractions were centrifuged at 16,000xg at 4°C for 10 min, and protein concentration was determined using the BCA protein assay kit.

### Flow Cytometry

Wild-type and ATGL-overexpressing MEFs or AML12s were cultured in 6-well plates with DMSO or etoposide (30 µM) for up to 48 hours. Following, cells were lifted, centrifuged, and fixed in 200 μL of PBS and 100 μL of 4% PFA for 15 minutes at room temperature. Cells were then washed with FACS buffer (PBS, 1% BSA, 0.5 mM EDTA) and stained with BODIPY 493/503 (2 µM) for 30 minutes at room temperature. Finally, they were washed, centrifuged, and resuspended in FACS buffer for flow cytometry data acquisition. Flow cytometry data were acquired with a FACSymphony A3 flow cytometer (BD Biosciences) and analyzed with FlowJo software (BD Biosciences). Relative lipid accumulation was calculated as the mean fluorescence intensity (MFI) of BODIPY in single cells.

