## Supplemental for "ATGL-catalyzed lipid catabolism promotes DNA double-strand break repair" (from another version of this preprint: version 1, 2026-04-04)

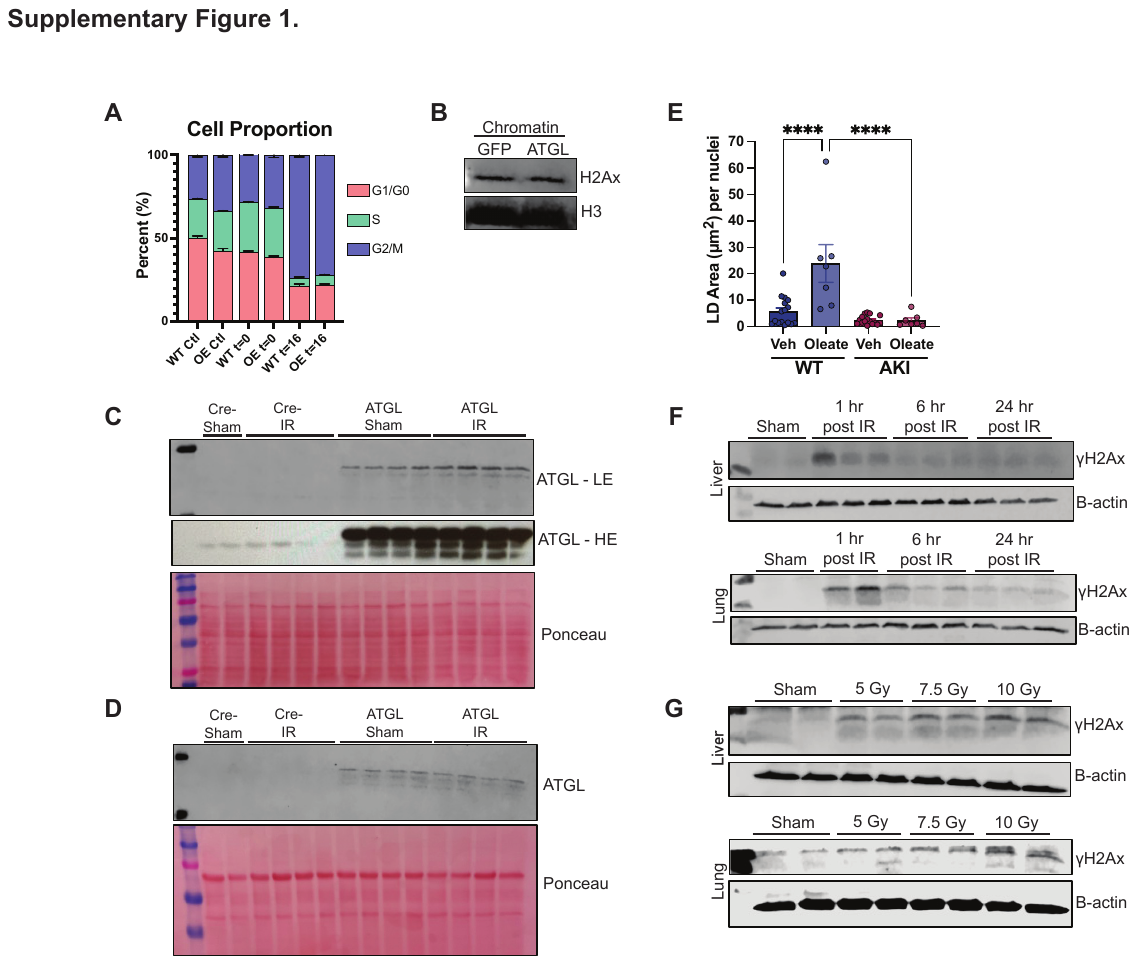

**Supplemental Figure 1. A)** Propidium iodide staining of control and ATGL overexpressing cells with etoposide treatment. “WT ctl”= null virus, no etoposide. “OE Ctl” = ATGL overexpression, no etoposide. “t=0” =cells harvested acutely after 3 hr in etoposide. “t=16” = cells harvested 16 hours after etoposide removal. **B)** Total H2Ax measurements performed on chromatin fractions of ATGL overexpressing or GFP control AML12 cells. **C)** γH2Ax dose response in WT mice at indicated IR doses. **D)** γH2Ax time course, harvested at indicated times post-IR. **E,F)** Validation of ATGL overexpression in E2A Cre-AKI liver (E) and lung (F). **G)** Primary MEFs were treated with 100 µM oleate overnight to induce LD accumulation, followed by one hour of oleate washout. Cells were stained with BODIPY 493/503 and imaged.

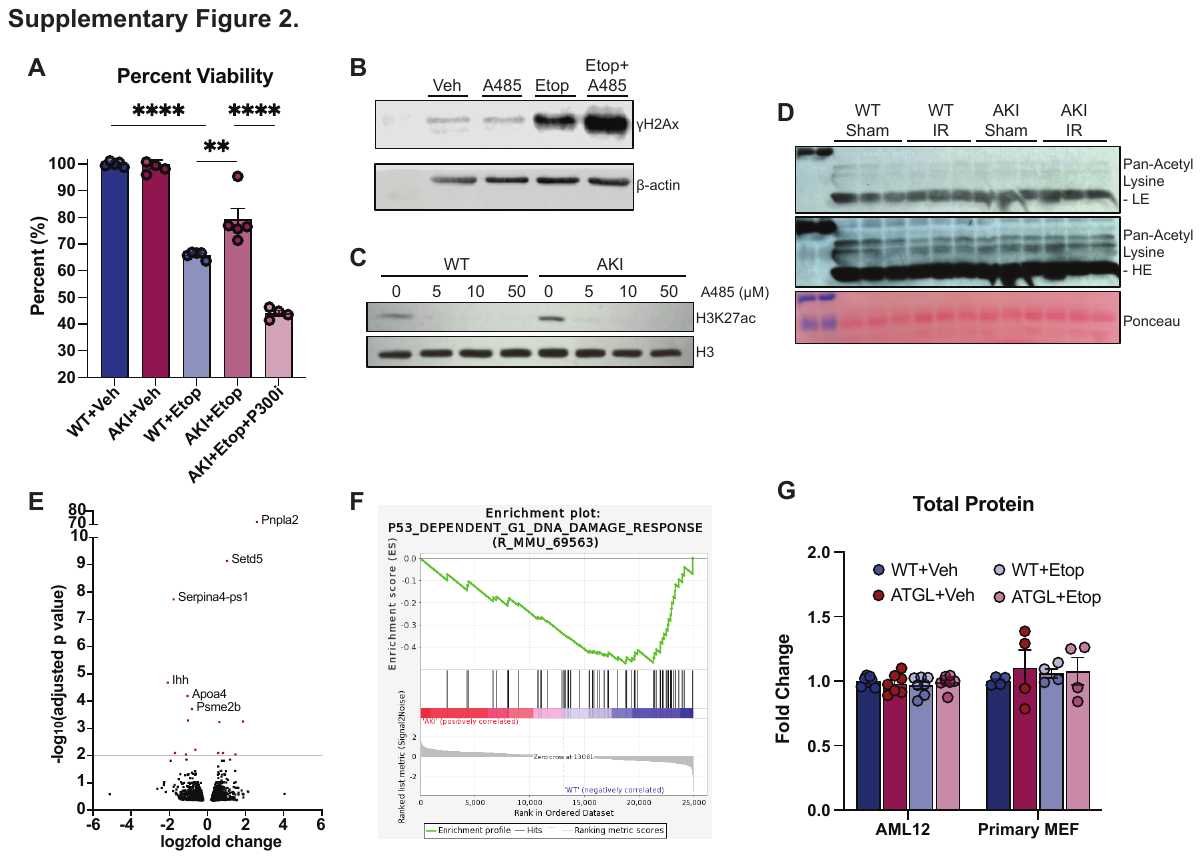

**Supplementary Figure 2. A)** Volcano plot of bulk RNA seq data. Significant genes (FDR <0.01, grey line) are shown in red. **B)** GSEA plot of genes enriched in AKI versus WT mice using Reactome predefined gene set R-MMU-69563 for p53-dependent DNA damage response.  **C)** Primary MEFs were pretreated with the P300 inhibitor A485 for 12 hours, followed by etoposide + A485 for 60 hours. Viability measured with Alamar Blue. Statistics: one-way ANOVA with Tukey’s post hoc test. **=p<0.005, ****=p<0.0001. **D)** Primary MEFs were pre-treated for 12 hours with 10 µM A485 followed by 6 hours of etoposide + A485 treatment. **E)** Primary MEFs were treated with denoted concentrations of A485 for 12 hours. **F)** Histone extract from WT and AKI mouse livers blotted for pan-acetyl lysine. **G)** Total protein measured by BCA from WT and ATGL overexpressing cells treated with etoposide.

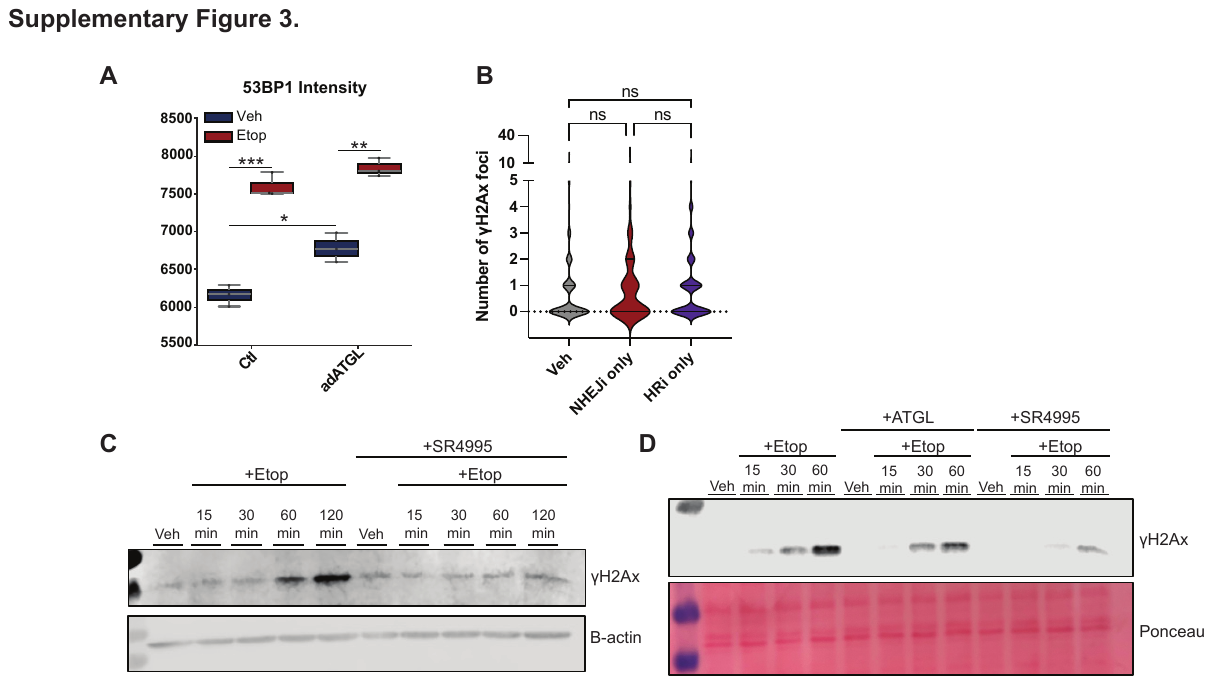

**Supplementary Figure 3. A)** Total nuclear intensity of 53BP1. AML12 cells stably expressing the mApple-53BP1 plasmid were transduced with adATGL. Cells were imaged after 3 hours of etoposide and 24 hours of recovery. **B)** Primary MEFs were treated with an NHEJ inhibitor (Nu7441, 10 µM) or HR inhibitor (RI-1, 20 µM) for 12 hours and stained for γH2Ax. Quantification of γH2Ax foci is graphed. Statistics: One-way ANOVA. **C)** IMR90 cells were pre-treated with 10 µM SR-4995 for 24 hours. Cells were treated with etoposide and were harvested at the indicated times. **D)** AML12 cells were transduced with ATGL or null virus control, or pre-treated with 10 µM SR-4995 for 24 hours. Cells were treated with etoposide and were harvested at the indicated times.
